# Process-based modelling of microbial community dynamics in the human colon

**DOI:** 10.1101/2022.07.04.498646

**Authors:** Helen Kettle, Petra Louis, Harry J. Flint

## Abstract

The human colon contains a dynamic microbial community whose composition has important implications for human health. In this work we build a process-based model of the colonic microbial ecosystem and compare with general empirical observations and the results of in-vivo experiments. Based our previous work (Kettle et al., 2015), the microbial model consists of 10 microbial functional groups, 4 substrates and 10 metabolites; to this we add the interaction with a human host to give simulations of the in-situ colonic microbial ecosystem. This model incorporates absorption of short chain fatty acids (SCFA) and water by the host through the gut wall, variations in incoming dietary substrates (in the form of “meals” whose composition varies in time), bowel movements, feedback on microbial growth from changes in pH resulting from SCFA production, and multiple compartments to represent the proximal, transverse and distal colon. We verify our model against a number of observed criteria, e.g. total SCFA concentrations, SCFA ratios, mass of bowel movements, pH and water absorption over the transit time; and then run simulations investigating the effect of colonic transit time, and the composition and amount of indigestible carbohydrate in the host diet, which we compare with in-vivo studies. Gut microbiota are highly complex and poory understood yet our work shows that it is nevertheless possible to develop predictive models of the key components of the dynamics of this ecological system. The code is available as an R package (microPopGut) to aid future research.

**Author Summary:** Kettle wrote the model code and led the writing of the manuscript. Louis and Flint both contributed to writing the manuscript and all aspects of microbiolgy. All authors contributed critically to the drafts and gave final approval for publication.

## Introduction

The human colon harbours a dense and diverse community of microbiota whose interactions with the host can have a profound effect on human health (e.g. Ros-Covin et al. (2016), Morrison and Preston (2016)). Due to the location of this community within its host, data collection and experimentation are problematic. Information on this system must come from volunteer experiments in which diet and stool samples are monitored or from laboratory experiments using the microbes found in stool samples. Another approach is to put current knowledge into a mathematical framework and run simulations of the system to test our understanding and identify knowledge gaps. To this end a number of mathematical models of this system have been developed - e.g. Cremer et al. (2016), Cremer et al. (2017), Munoz-Tamayo et al. (2010), Smith et al. (2021), Moorthy et al. (2015).

When developing a model, a number of assumptions about the system are made in order to reduce complexity/dimensionality so that the model is easier to parameterise, run and analyse. Some modellers choose to reduce the microbial complexity and focus on the physics of the gut (e.g. Cremer et al. (2016), Cremer et al. (2017)), some try to achieve a balance of both (e.g. Munoz-Tamayo et al. (2010)) and some choose to develop the microbial community (e.g. Smith et al. (2021)). The model described here focuses on the microbial community dynamics and on interactions with the host, with a fairly simple model of the colon. We include the simulation of ‘meals’ (of random composition and size) arriving at the colon and look at the effects of bowel movements, both of which, as far as we are aware, have not been previously incorporated into such models. Having developed a complex model of human gut microbiota in a fermentor system (Kettle et al., 2015), and publicly available software (microPop - an R package for modelling microbial communities (Kettle et al., 2018)) we now incorporate this 10-group microbial ecosystem model (Table 1) into a model of the human gut in order to simulate the effects of diet and host on the microbial composition and subsequent short chain fatty acid (SCFA) production.

**Table 1:** Microbial functional groups included in the model (and the R package microPop (Kettle et al., 2018)) and described by Kettle et al. (2015). Users should be aware that the parameter values given in the data frames in the software will almost certainly change with increasing knowledge of gut microbiota and in some cases are simply a “best guess”.

| microPop Name | Abbr. | Description | Examples |
| --- | --- | --- | --- |
| Bacteroides | B | Acetate-propionate-succinate group | *Bacteroides* spp., *Prevotella* spp., *Akkermansia muciniphila* (Verrucomicrobia) |
| NoButyStarchDeg | NBSD | Non-butyrate-forming starch degraders | *Ruminococcaceae* related to *Ruminococcus bromii*. Also includes certain *Lachnospiraceae* |
| NoButyFibreDeg | NBFD | Non-butyrate-forming fibre degraders | *Ruminococcaceae* related to *Ruminococcus albus*, *Ruminococcus flavefaciens*. Also includes certain *Lachnospiraceae* |
| LactateProducers | LP | Lactate producers | *Actinobacteria*, especially *Bifidobacterium* spp., *Collinsella aerofaciens* |
| ButyrateProducers1 | BP1 | Butyrate Producers | *Lachnospiraceae* related to *Eubacterium rectale*, *Roseburia* spp. |
| ButyrateProducers2 | BP2 | Butyrate Producers | Certain *Ruminococcaceae*, in particular *Faecalibacterium prausnitzii* |
| PropionateProducers | PP | Propionate producers | *Veillonellaceae* e.g. *Veillonella* spp., *Megasphaera elsdenii* |
| ButyrateProducers3 | BP3 | Butyrate Producers | *Lachnospiraceae* related to *Eubacterium hallii*, *Anaerostipes* spp. |
| Acetogens | A | Acetate Producers | Certain *Lachnospiraceae*, e.g. *Blautia hydrogenotrophica* |
| Methanogens | M | Methanogenic archaea | *Methanobrevibacter smithii* |

Approximately 95% of the SCFA produced by the microbes during growth are absorbed by the host through the gut wall and it is the ratio of the 3 main SCFAs (acetate, butyrate and propionate) which is known to have a significant effect on human health. Thus, we prioritise information on the values of these ratios in our model verification. Similarly approximately 90% of the water flowing into the colon is absorbed. Changes in the volume of water have a significant effect on the concentration of the molecules in the colon which in turn affects pH which then affects microbial growth, all of which are included in our model.

Due to its shape within the body, the colon is commonly divided into 3 different regions - the proximal, transverse and distal sections running from beginning to end (Fig. 1A and B). The availability of substrate, microbial growth and hence pH vary along the colon, therefore, although our model is not spatial we simulate these three regions explicitly, with flow from one to another. Furthermore, as well as incorporating varying substrate inflow in the form of meals we also add in the release of mucins along the length of the colon which can be microbially broken down to release proteins and carbohydrates, allowing for further microbial growth away from the beginning of the colon where the substrates enter. A graphical summary of the model is shown in Fig. 2.

**Figure 1:**
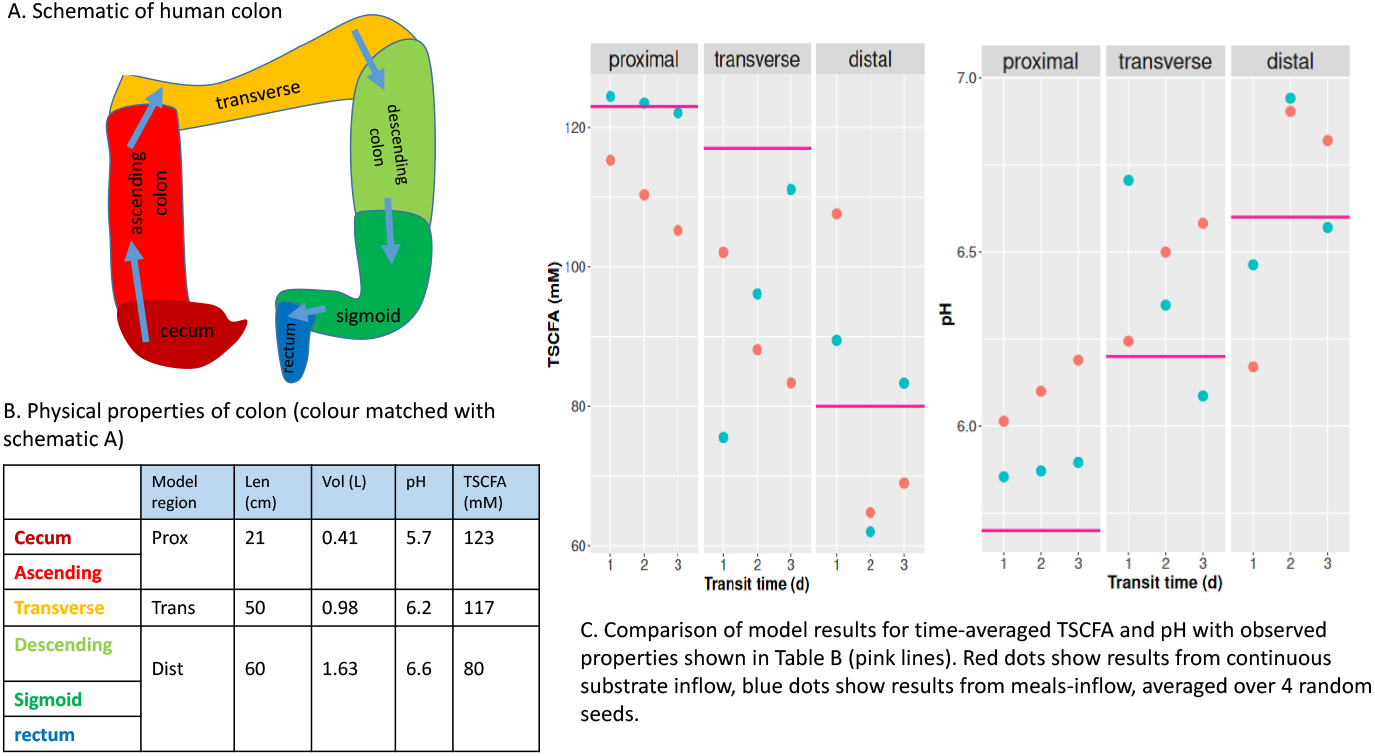
Colon schematic plus table of typical values for physical properties (length, volume, pH and TSCFA) and plots of summarised model simulations for average TSCFA and pH for comparison with typical values.

**Figure 2:**
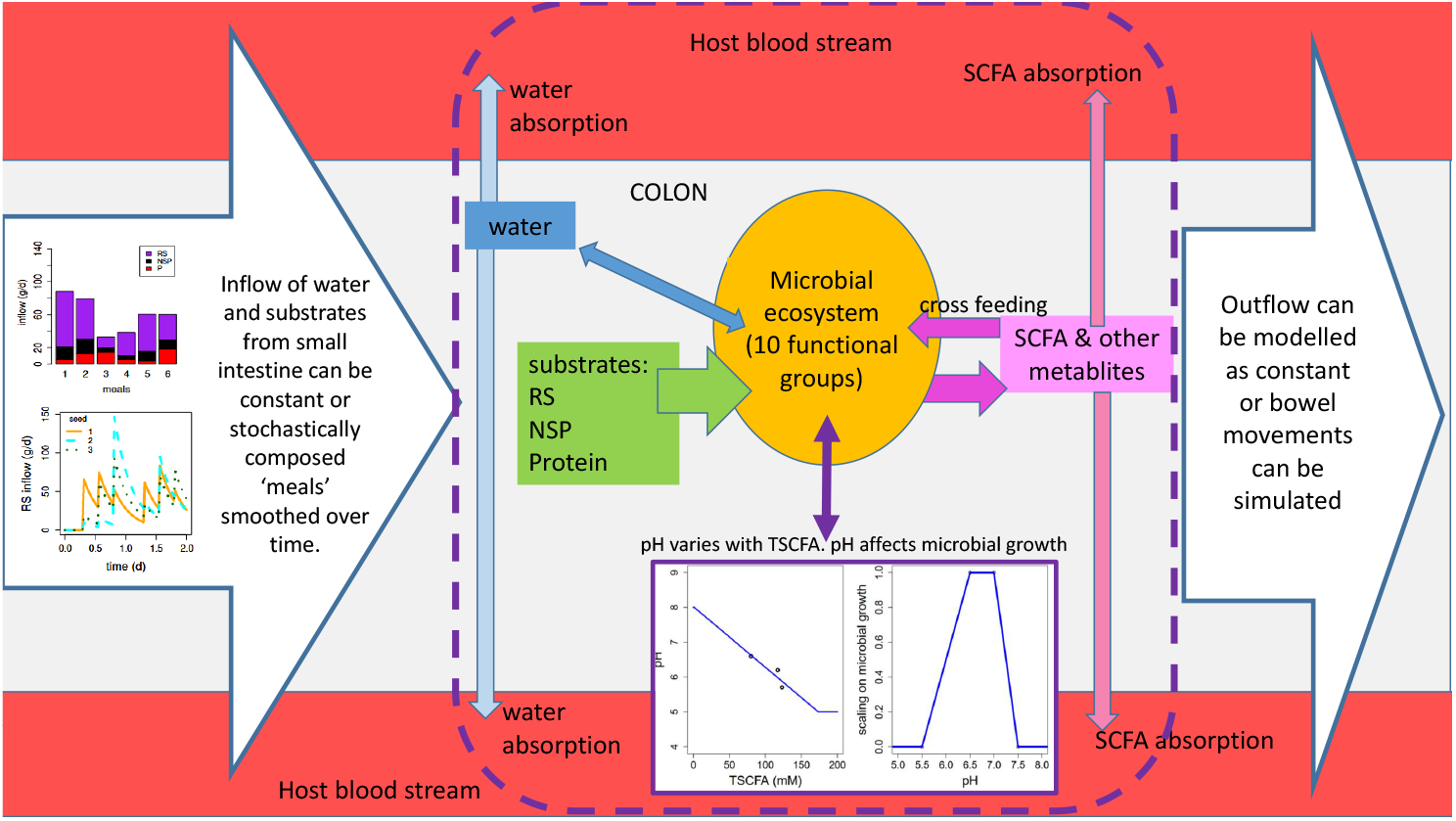
Model system with the microbial ecosystem comprising 10 microbial functional groups (Table 1) which consume substrates (RS, NSP and protein) and water. The microbes produce metabolites some of which are consumed by other MFGs (‘cross-feeding’). SCFA and water are absorbed through the colon wall (at a different specific rates). The system shown within the dashed line is repeated in each of the modelled regions of the colon (proximal, transerve, and distal) with the contents of the previous region, flowing into the next. The first compartment (proximal) has inflow from the small intestine - this can be constant inflow or simulated meals whose composition varies randomly in time. The third model compartment (distal) has outflow to stool which can be constant or evacuation via bowel movements can be simulated. pH varies with the TSCFA concentration and affects the rate of microbial growth differently for each MFG.

We use the following criteria to verify our model captures the main features established for the system:

1. Total SCFA (TSCFA) concentration in the proximal, transverse and distal compartments should be around 123, 117 and 80 mM respectively according to sudden death human autopsies (Cummings et al., 1987)
2. Acetate:Propionate:Butyrate ratios are similar (around 3:1:1) in all regions of the colon and around 60:20:20 mM (Cummings et al., 1987)
3. Over 95% of SCFA are absorbed by the host (Topping and Clifton, 2001)
4. Approx. 90% of incoming water is absorbed by the host (Phillips and Giller, 1973)
5. pH in the proximal, transverse and distal compartments should be around 5.7, 6.2 and 6.6 respectively (Cummings et al., 1987)
6. Normal daily fecal output in Britain is 100-200 g d^*−*1^ of which 25-50 g is solid matter (i.e. 50-175 g d^*−*1^ is water). Bacteria make up about 55% of the solid matter i.e. 14-28 g d^*−*1^ of microbes emitted (Stephen and Cummings, 1980).
7. TSCFA concentration decreases with transit time (Lewis and Heaton, 1997)

After model verification we examine the effects of including meals, bowel movements and fixed/varying pH into the model. We then use the model to look at how carbohydrate composition (based on the fractions of resistant starch (RS) and non-starch polysaccharides (NSP)) and total carbohydrate affect the microbial community and SCFA composition. The simulations are then compared with in-vivo data from human volunteer experiments.

Although gut microbiota are highly complex and not fully understood, here we show that it is nonetheless possible to develop predictive models of key components of this ecological system. Our results show promise and we believe this model represents a significant step forward in this field. We refer to the model as “microPopGut” and to aid future research the code is available as an R-package on github (https://github.com/HelenKettle/microPopGut) and instructions on how to use the package are given in the supplementary file ‘gettingStartedWithMicroPopGut.pdf’.

## Results

### Standard Model

Having established the default model settings and parameter values which give the best fit to our criteria (see Table 3 for colon parameters and Supp. Info. (section 3) for microbial group parameters) we then investigate the effects of different model configurations, e.g. with/without bowel movements, meals and variable pH, for a range of transit times. Simulations with meals have a random component therefore the model is run for a number of different starting seed values. Due to the random fluctuations these simulations will not reach steady state therefore the summary values are taken as the mean from day 7 (to remove the effect of the initial conditions) to the end of the simulation (28 days) and are averaged over multiple seeds.

Table 2 gives summary results of the model simulations without bowel movements but with varying pH for each bowel region. Fig. 3 shows results from more simulations but for the distal colon only. Fig. 3a shows that although bowel movements make a difference to the total biomass and the TSCFA they do not have a large effect on the community composition or the SCFA ratios. Thus in the interests of model simplicity we decide to not include bowel movements in later simulations. However, varying pH with TSCFA can be seen to make a very large difference to the microbial community (Fig. 3b) and also improves the SCFA ratios with respect to our verification criteria. The addition of meals makes a significant difference which increases with increasing transit time (Fig. 3c). In Fig. 4 the time series output from the model shows how the meals-inflow allows the community to experience large shifts over time (on a much longer time scale than the variations in the input), as opposed to the fixed state approached using a constant substrate inflow.

**Table 2:**
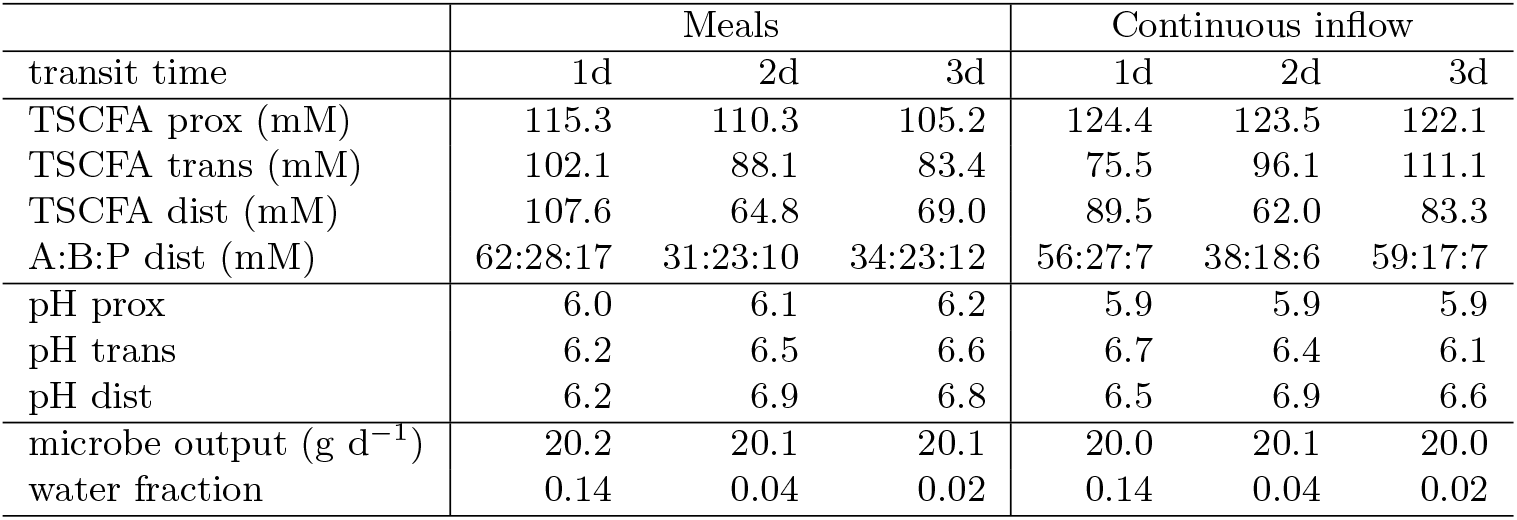
Summary of model results (for comparison with our list of criteria) for 3 different transit times, with meals or continuous inflow and with pH varying with TSCFA. Microbe output is the mass of microbes leaving the colon per day and the water fraction is amount of water leaving the colon per day divided by the amount entering. All simulations were run for 28 days and the results shown are the average over days 7-28. The results for the simulations with meals are averaged over 4 random seeds. ‘A:B:P dist’ refers to the Acetate:Butyrate:Propionate ratio (mM) in the distal colon.

| transit time | Meals |  |  | Continuous inflow |  |  |
| --- | --- | --- | --- | --- | --- | --- |
|  | 1d | 2d | 3d | 1d | 2d | 3d |
| TSCFA prox (mM) | 115.3 | 110.3 | 105.2 | 124.4 | 123.5 | 122.1 |
| TSCFA trans (mM) | 102.1 | 88.1 | 83.4 | 75.5 | 96.1 | 111.1 |
| TSCFA dist (mM) | 107.6 | 64.8 | 69.0 | 89.5 | 62.0 | 83.3 |
| A:B:P dist (mM) | 62:28:17 | 31:23:10 | 34:23:12 | 56:27:7 | 38:18:6 | 59:17:7 |
| pH prox | 6.0 | 6.1 | 6.2 | 5.9 | 5.9 | 5.9 |
| pH trans | 6.2 | 6.5 | 6.6 | 6.7 | 6.4 | 6.1 |
| pH dist | 6.2 | 6.9 | 6.8 | 6.5 | 6.9 | 6.6 |
| microbe output ($\text{g d}^{-1}$) | 20.2 | 20.1 | 20.1 | 20.0 | 20.1 | 20.0 |
| water fraction | 0.14 | 0.04 | 0.02 | 0.14 | 0.04 | 0.02 |

**Figure 3:**
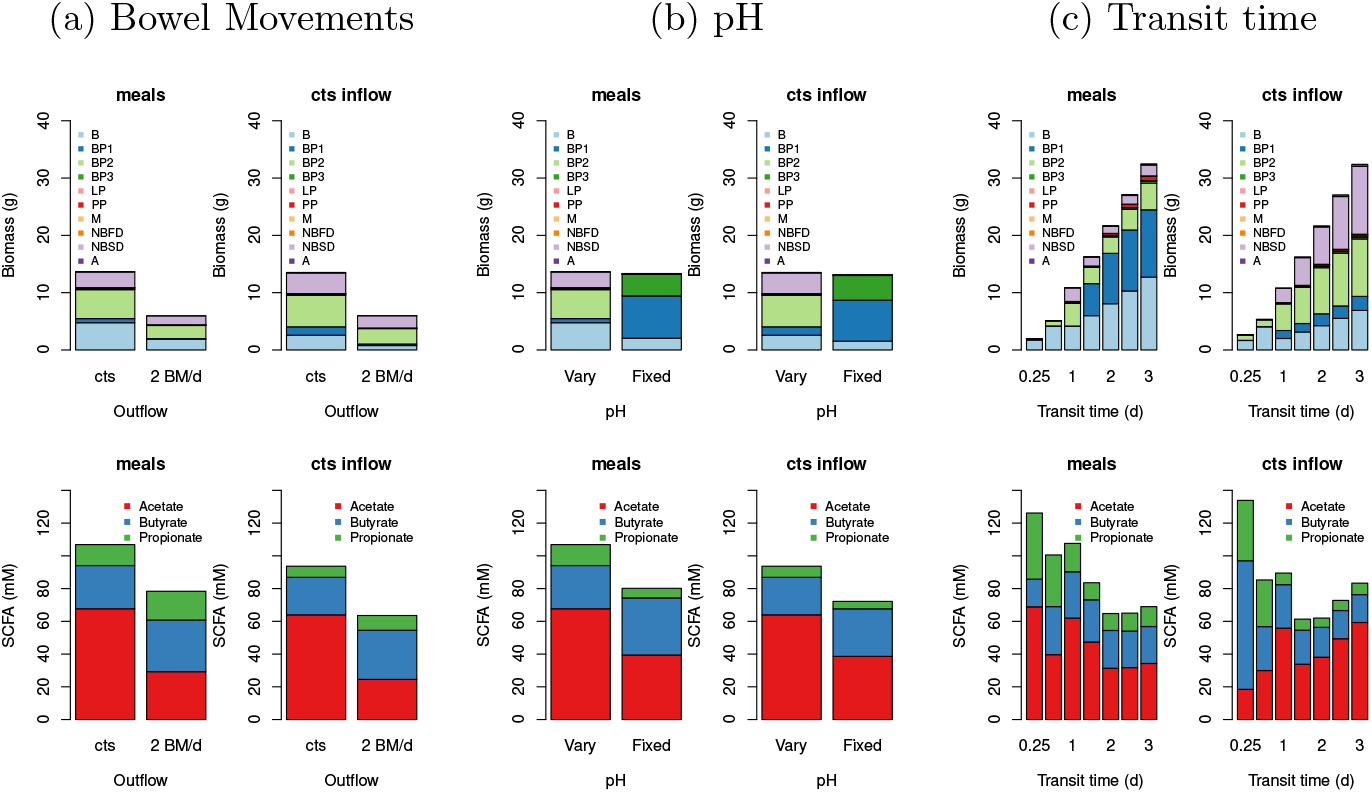
Summary results (averaged over days 7-28 and over random seeds) for the distal compartment for continuous inflow or fluctuating inflow (i.e. ‘meals’) for continuous outflow from colon or for 2 bowel movements per day (‘2 BM/d’). The RS fraction is 0.78 (i.e. 78% of the dietary carbohyrate is resistant starch and 22% is NSP) and the transit time is 0.93 d for a), 1.25 d for b) and at 0.25, 0.5, 1, 1.5, 2, 2.5 and 3 days for c). The top row shows the biomass of each group, the bottom row shows the SCFA.

**Figure 4:**
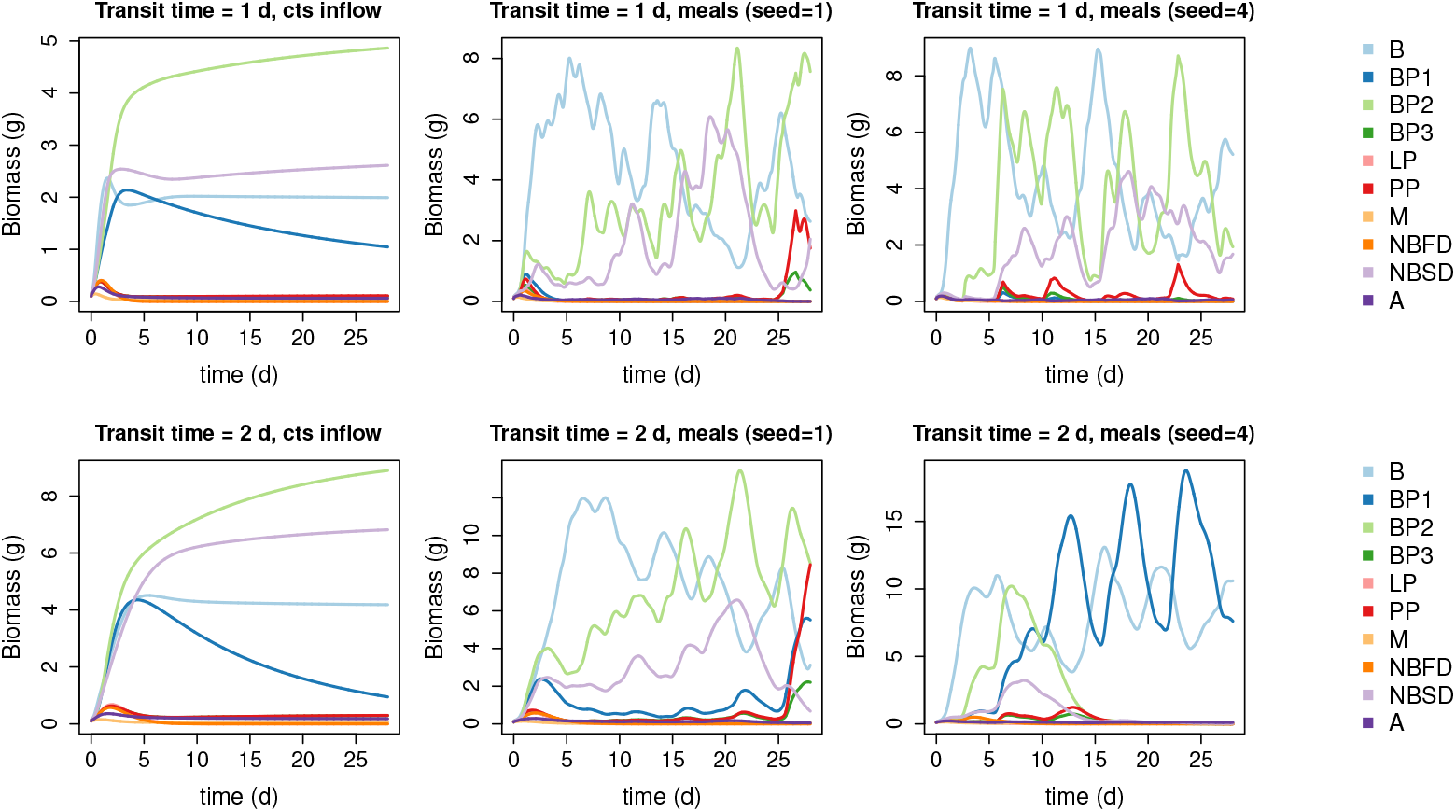
Simulation results for the distal compartment for continuous inflow (first plot on each row) or fluctuating inflow (i.e. ‘meals’) for transit times of 1 d (top row) and 2 d (bottom row) and for 2 random seeds. Modelled pH varies with TSCFA and the RS fraction is 0.78. There are no bowel movements (i.e. outflow is continuous). See Table 1 for microbial groups.

Fig. 1C shows the average pH and TSCFA for the proximal, transverse and distal compartments. It can be seen that blue (meals) and red (continuous inflow) dots show the same basic trends. The decrease in TSCFA with transit time has been shown experimentally (Lewis and Heaton, 1997); in section 2 of the Supp. Info. we suggest a mathematical explanation for this based on the supposition that the specific rate of absorption of water through the gut wall is slower than that for SCFA.

Regarding Table 2, for some criteria, e.g. pH, the continuous inflow setting gives results closer to our verification values, but in other cases, e.g. A:B:P in distal colon, simulating meals gives closer results. Note that we consider a transit time of 1 day the most typical of the three transit times, and the one that should be compared with our verification criteria, the others are included to show the variation in results. Ideally TSCFA should be 123, 117 and 80 mM for prox., trans., dist. but the best match we have to this is for a 3 d transit time and continuous inflow. This is most likely due to the fact that our model has fixed rates of specific absorption of SCFA and water throughout the colon. However, our TSCFA values are within a reasonable range and display the general trend of decreasing TSCFA from the proximal to distal colon. The microbe output, i.e. the outflow of fecal microbes is steady at around 20 g d^*−*1^ in all cases which fits well in the verification range (14-28 g d^*−*1^). The water fraction is the ratio of the rate of fecal water over the rate of water flowing into the colon, since 90% of water is absorbed this should be 0.1. This is approximately correct for our 1 d simulations (0.14) but, as expected, when transit time increases this decreases significantly. In summary, comparing these simulation results with our list of model verification criteria shows that in general our model is fit for purpose, and that the inclusion of meals-inflow and varying pH improve our simulations.

### Model Experiments

We now use our model to simulate two scenarios – firstly, the effects of decreasing total carbohydrate intake and secondly, the effects of changing carbohydrate composition (whilst keeping total intake fixed) on the microbial commnuity and associated SCFA production. Comparing our simulations with data from human volunteer experiments is not straightforward since in order to run our model, ingested food must be translated to substrates reaching the colon. This is problematic due to unknown water consumption and transit times and uncertainties associated with the absorption rates of the ingested carbohydrate and protein higher up the digestive tract. Thus we do not attempt to reproduce human experiments but rather we run simulations based on variations to our standard model set up and then compare our results qualitatively with available data.

#### Effects of total dietary carbohydrate

In this model experiment we investigate the effects of decreasing carbohydrate on the microbial community. Here we compare our results qualtitatively with the human dietary study of Duncan et al. (2007) which explored the impacts of carefully controlled decreases in carbohydrate intake upon weight loss and microbial fermentation products in obese subjects using 3 diets – a maintenance (M) diet, a high protein, moderate carbohydrate diet (HPMC) and a high protein, low carbohydrate diet (HPLC) (see Fig. 5 for details). This is of course, the composition for ingested food, which is not easily translated into substrate concentrations entering the colon. However, we can look at the general trends in SCFA and microbial composition with changing colonic carbohydrate intake rate. Thus, in these model experiments we keep protein inflow to the colon at 10 g d^*−*1^ (our default value) and then increase inflowing carbohydrate from 10 g d^*−*1^ to 60 g d^*−*1^ in 10 g d^*−*1^ intervals. To include the effects of different carbohydrate composition we run the model for an resistant starch (RS) fraction of either 0.2 or 0.78 (the default value), with non-starch polysaccharides (NSP) making up the remaining carbohydrate in each case. Although subject to large uncertainties, we estimate the RS fractions for the Duncan et al. (2007) experiments of 0-0.6 (M diet), 0-0.68 (HPMC) and 0-0.12 (HPLC) (based on RS is 0–20% of ingested starch (Capuano et al., 2018) and bio-available NSP is 75% of ingested NSP (Slavin et al., 1981)). Due to the low fibre nature of many of these simulations we run the model with a slightly longer transit time of 1.5 d and for both continuous inflow and meals.

**Figure 5:**
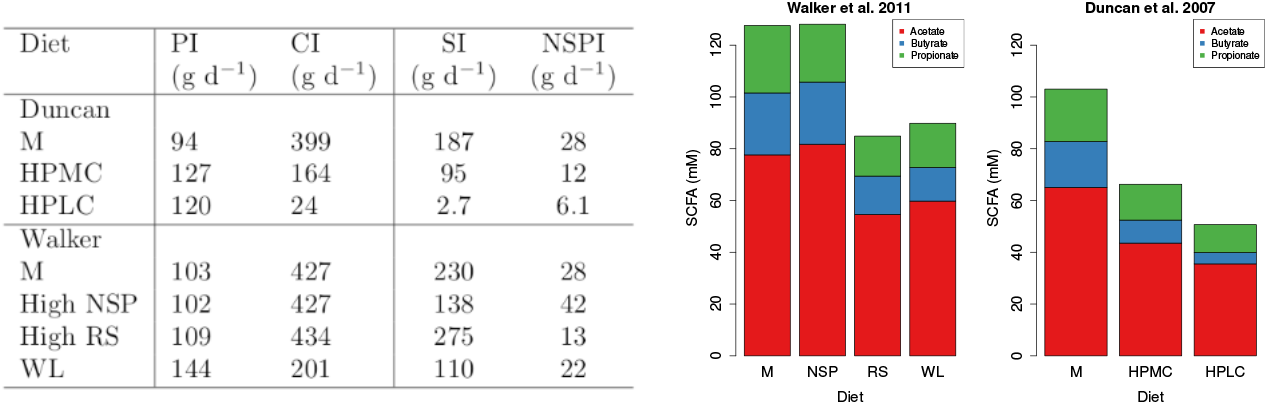
Table on left shows the dietary intake for two human studies (Duncan et al. (2007) and Walker et al. (2011). PI, CI, SI and NSPI refer to ingested dietary protein, carbohydrate, starch and NSP. Note, starch value for the high RS diet in the Walker et al. (2011) study included 26 g commercial RS. Bar plots show SCFA data from these studies.

Fig. 6 shows the SCFA results from our model experiment and Fig. 5 shows the results from the in vivo experiment. It is very clear, from both the model and in vivo results that the proportion of butyrate increases as the amount of carbohydrate in the diet increases. Furthermore, both model and in vivo results show an increase in TSCFA with carbohydrate intake rate. Since Duncan et al. (2007) also look at the relationship between butyrate concentration and grams of carbohydrate eaten per day we plot butyrate against carbohydrate entering the colon (Fig. 7) to compare with their Fig. 1. In both cases, butyrate concentration increases with incoming carbohydrate. Furthermore, as seen in both the model and the data, the percentage of butyrate increases with carbohydrate intake (Fig. 7).

**Figure 6:**
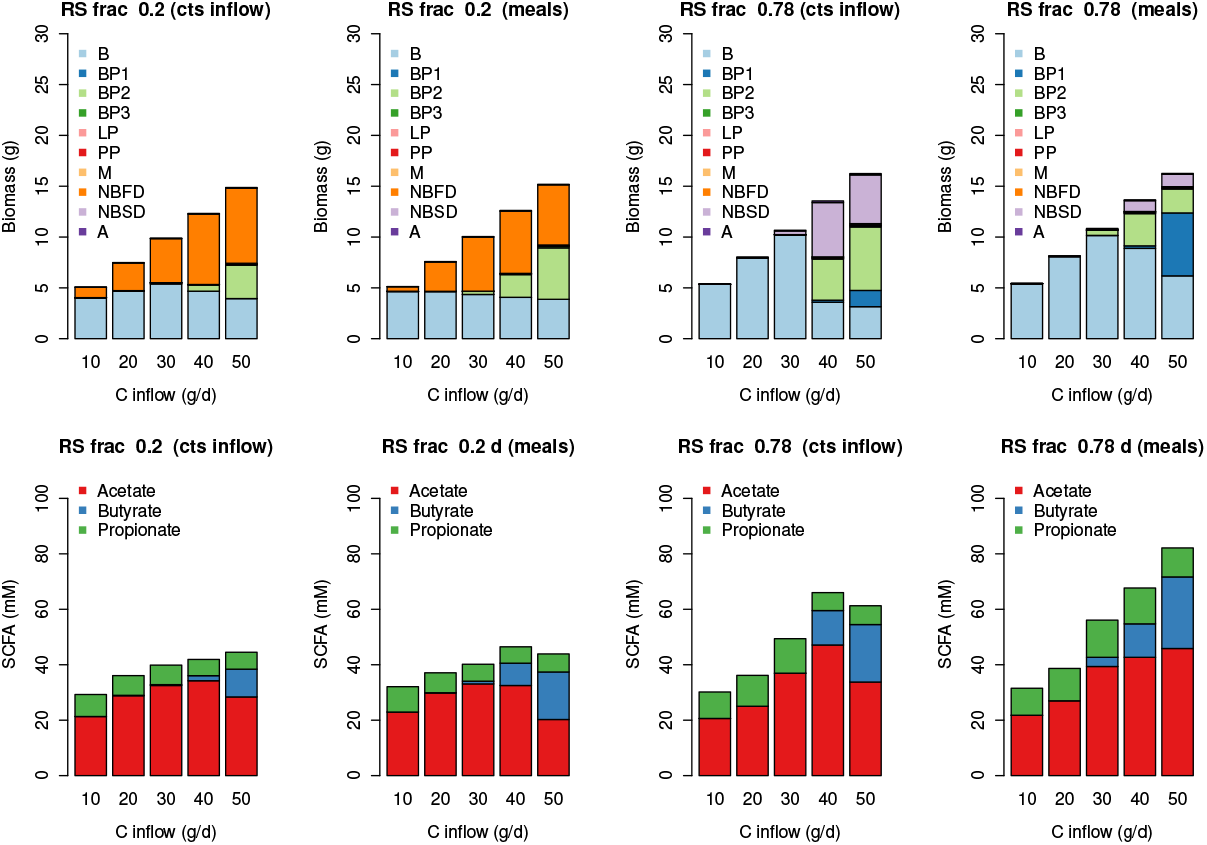
Simulated Biomass and SCFA results for increasing carbohydrate inflow. Simulations are run with continuous substrate inflow (cts) and with ’meals’ for a transit time of 1.5 days. The results are the average over the last 3 weeks of a 28 day simulation and ’meals’ is the average over 4 stochastically-generated simulations.

**Figure 7:**
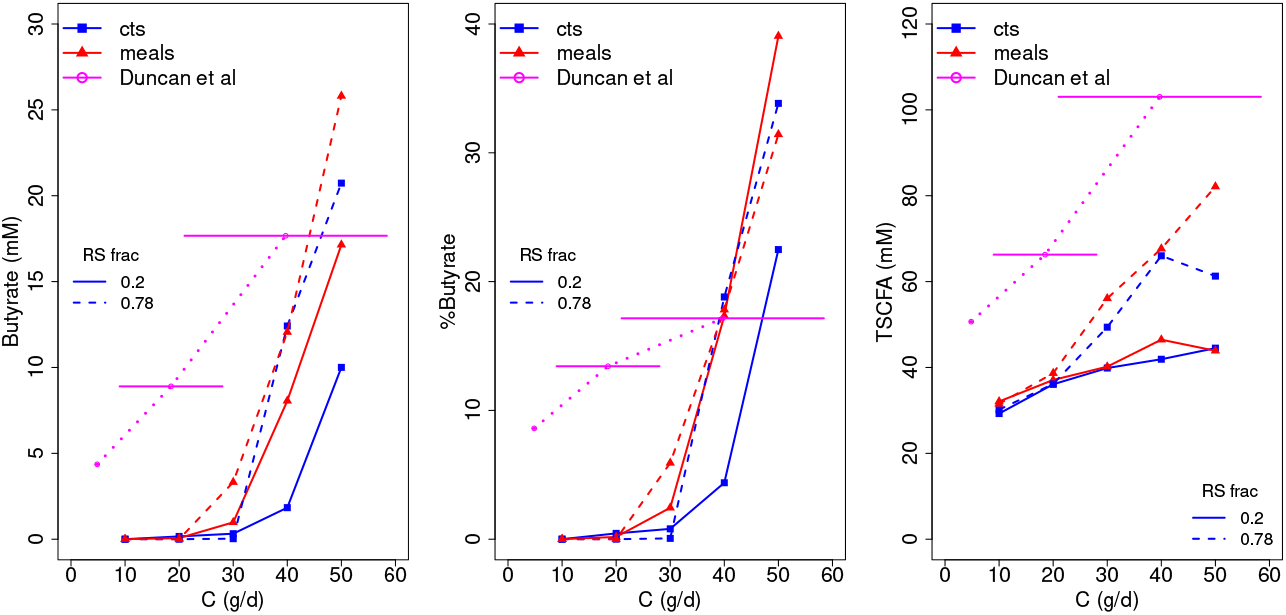
Plot of modelled butyrate, %butyrate and TSCFA against grams of carbohydrate entering the colon each day. Data from Duncan et al. (2007) is shown in magenta - due to uncertainties in converting ingested starch to RS entering the colon there are large error bars on the amount of C (g/d). Error bars show C estimated by the sum of 75% of ingested NSP plus 0-20% of ingested starch.

In terms of microbial composition, Fig. 6 shows the results from our simulations are reasonably consistent across inflow type (meals or continuous), with B dominating at low carbohydrate intake. When the RS fraction is low (i.e. when carbohydrate is made up of 80% NSP) then NBFD increase with increased C intake. Whereas when C is mostly RS then NBSD and BP1 increase with C. In both cases BP2 increase with increasing C intake.

#### Effects of carbohydrate composition

Here we use the model to simulate the effects of changing carbohydrate composition on the microbial community composition by changing the ratio of RS to NSP whilst keeping the same amount of total incoming carbohydrate. Fig. 8 show a summary of the model results. Although there are differences between the continuous inflow/meals, and also for the different transit times (1 d and 3 d), the modelled trends are generally similar, showing a significant shift in community as the fraction of RS increases, an increase in TSCFA and changes in the SCFA ratios. We compare our results with a human dietary study (Walker et al. (2011), Salonen et al. (2014) and references therein) examining the impact of switching the major type of carbohydrate from wheat bran (NSP) to resistant starch. Volunteers were provided successively with a maintenance diet, diets high in RS or NSPs and a reduced carbohydrate weight loss (WL) diet, over 10 weeks (Fig. 5).

**Figure 8:**
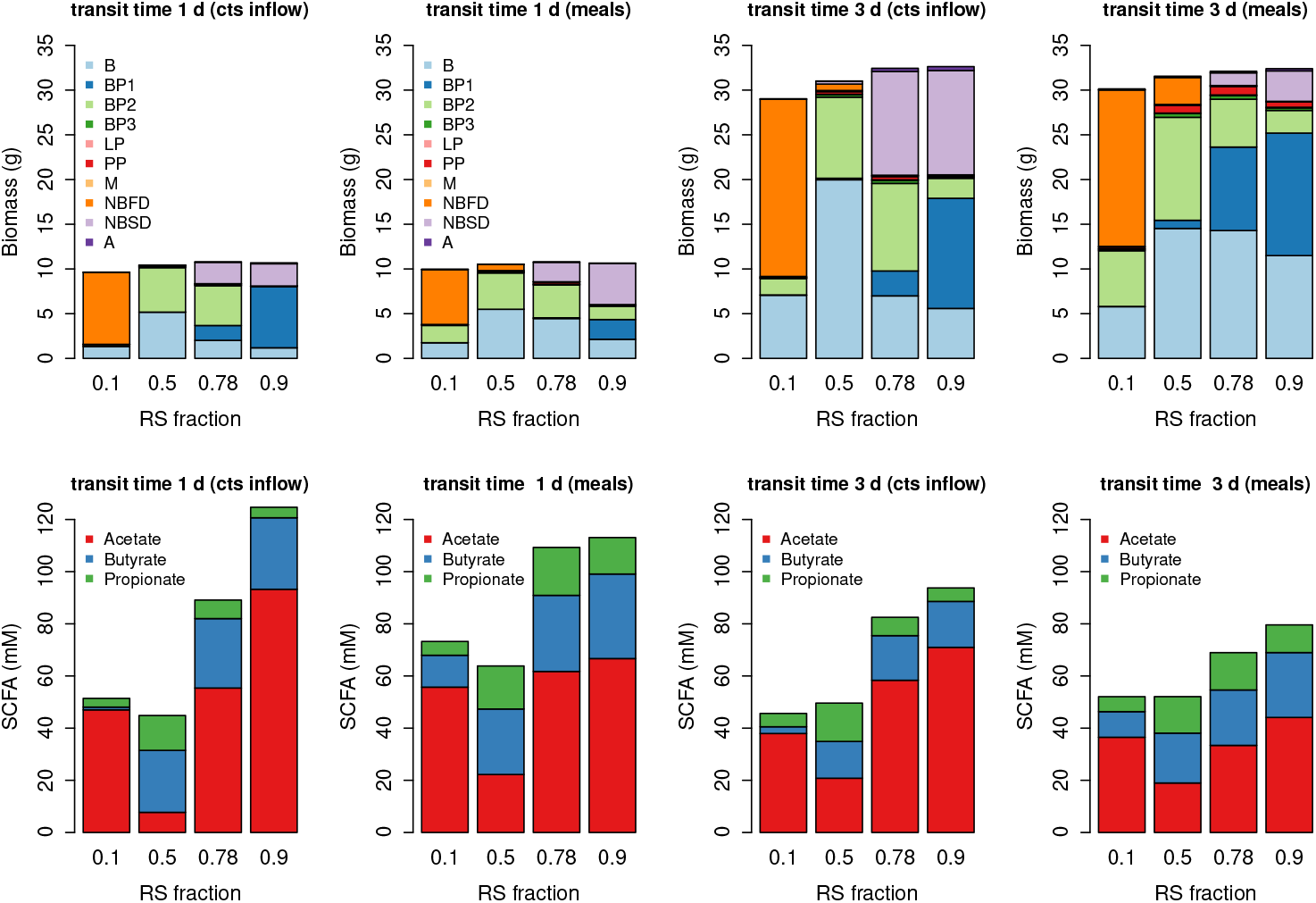
Biomass and SCFA results for changing the RS fraction of inflowing carbohydrate with continuous substrate inflow (‘cts inflow’) and with ‘meals’. Protein and carbohydrate inflow are 10 and 50 g d^*−*1^ respectively. The results are the average over the last 3 weeks of a 28 day simulation and ‘meals’ is the average over 4 stochastically-generated simulations.

There are very large discrepancies between the SCFA predicted by our model (Fig. 8) and the measured SCFA data (Fig. 5). Our model predicts an increase in TSCFA as proportion of RS increases whereas total fecal SCFA were significantly lower for the RS and WL diets compared to the other two diets (in which NSP is higher). One possible explanation is that fermentation of RS occurs in more proximal regions of the colon compared with NSP fibre fermentation, such that there is greater absorption of the SCFA products. A second possibility, also likely, is that transit times were longer for the RS diet than for the NSP diet, which we predict would result in decreased SCFA concentrations.In our model the effect of the RS fraction on TSCFA is greater than the effect of transit time so we do not see this in Fig. 8.

The human study also included detailed compositional analysis of the fecal microbiota (Walker et al. (2011), Salonen et al. (2014)) that revealed specific responses mainly by different groups of Firmicutes bacteria to the RS and NSP diets. This information was particularly important for the phylogenetic assignments to the functional groups used here and previously in the model of Kettle et al. (2015). Our modelling predicts striking shifts in the microbial community, especially involving the NBSD, NBFD and butyrate-producing groups, with changing proportions of RS and NSP fibre (Fig. 8). We should also note that in the volunteer experiments many bacterial species were not significantly altered by the RS-NSP switch in vivo (Walker et al., 2011) indicating that many may be generalists, able to switch quickly between energy sources.

## Discussion

The development of a complex model of the microbial community in the human colon, whose simulations compare well with data, represents a significant step forward in this field. Previous models have been based on simpler microbial models (e.g. Cremer et al. (2017), Munoz-Tamayo et al. (2010), Moorthy et al. (2015)), or have not shown such a good agreement with data (e.g. Smith et al. (2021)). Our previous complex model community consisted of 10 functional groups, but the model was designed only to simulate continuous culture conditions in a chemostat (Kettle et al., 2015). Translating this 10-group model into an in vivo setting has required introducing multiple gut compartments, and the absorption of water and SCFA, followed by comparison with generally observed characteristics of the system. We were then able to use this model to examine the predicted impact of changes in the amount and type of non-digestible carbohydrate (fibre) present in human diets upon concentrations of fermentation products (SCFA) in different gut compartments and in stool. At the same time, we predict the likely impact of dietary changes and variations in gut transit upon microbiota composition and fermentation products. The model must be regarded as work in progress particularly with respect to microbiota composition. Predictions can however become improved and refined as more information becomes available in time.

Assignments of microbial taxa to our ten functional groups were based initially on evidence from cultured isolates. These assignments have since been supported and greatly extended by analysis of genes diagnostic for different fermentation pathways within genomes and metagenomes (Reichardt et al., 2014) and by molecular detection of species enriched within the community by defined growth substrates in chemostat experiments (Duncan et al., 2016) and dietary intervention studies (Salonen et al., 2014). Nevertheless, these assignments inevitably remain provisional and incomplete and we do not claim that the model predictions can be made precise at a phylogenetic level. More emphasis is placed in our model on the prediction of metabolic outputs based on microbial transformations and interactions. While there is relatively little phylogenetic overlap for example between producers of propionate and butyrate (Reichardt et al. (2014), Louis and Flint (2017)) there are many cases where individual species are known to use multiple alternative substrates as energy sources, which complicates assignments. For this reason, more weight was given to fermentation pathways than to substrate preferences in defining the functional groups. It may well be worthwhile to increase the number of functional groups in the future. The large B group for example currently includes members of the Bacteroidetes phylum, but its characteristics are mainly based on well-studied members of the Bacteroides genus. We know that Prevotella is another highly abundant genus of Bacteroidetes in the human colon, but the two genera tend not to co-occur at high levels in the same individuals (Wu et al. (2011), Chung et al. (2020)). Less is known about human colonic Prevotella, for which there are relatively few cultured representatives, making it premature to create a separate grouping, but this would clearly be desirable in the future as their prevalence is reported to affect health and responses to dietary intervention.

The parameter values for the microbial groups used in our model are from the intrinsic data frames in the microPop package (the only changes are to LactateProducers). Although the work presented here did not attempt to fit particular parameters to data, as we focussed on expanding the scope of the model (i.e. changing the environment from fermentor to colon), these values are easy to alter, e.g. Wang et al. (2020) changed many of these parameters to achieve a better model fit to their data. Furthermore, it is also possible to include any number of strains (with varying parameter values) within each functional group in order to add more variation in outcome (see Kettle et al. (2015)) but we did not do this here in the interests of computational time. It should also be noted that the parameter values are highly uncertain in many cases and within each of our functional groups there will be large variability due to adaptation and evolution. Given this, we do not claim that the model response is necessarily representative of what may happen in an individual’s gut, rather it can be used as an aid to gain insight into the relative importance of the different processes we are currently aware of and potentially to highlight, those we are not.

In addition to this, it must be noted that the default diet chosen here with 10 g of protein and 50 g of carbohydrate fibre reaching the colon each day could be revised for any given population. However, converting from quantities of ingested food to substrate inflow to the colon is highly uncertain with large variations between studies, as well as technical issues with measuring this accurately. With more time, it would be interesting to investigate a larger range of typical diets but this was beyond the scope of the current work.

In summary, although performing reasonably well, the model has the potential to be considerably improved simply by altering the parameter values and existing settings, however, more fundamental changes such as those listed below could also be investigated in future work:

- Adding more functional groups or pathway switches in the existing functional groups. For example at present only the Bacteroides group can utilise protein but it is now known that some butyrate producers can also utilise amino acids (Louis and Flint, 2017)
- Our pH relation with TSCFA is very simplistic and could potentially be improved, although host secretions mean this is not necessarily straightforward.
- Currently we set the transit time for the colon and then this is split between the 3 model compartments based on their relative sizes. An interesting addition would be to alter transit time based on the composition of the various substrates entering the colon. For example, increasing residence time for high protein and/or low fibre diets. Due to variation in individual response this may need to include significant uncertainty ranges.
- Related to this is changing the absorption rate of water through the gut wall based on the diet, for example more water could remain in the gut on a high fibre diet.

To conclude, our model helps to explain some important, but poorly understood, relationships that have been reported from human dietary intervention studies, e.g. an increase in fecal total SCFA concentrations with faster gut transit. Gut transit is also shown to have potentially important consequences for microbiota composition and gut metabolism. In addition, the model confirms that the amount and type of non-digestible carbohydrate in the diet has the potential to cause major changes in microbiota composition. The nature of such changes is, however, predicted to be influenced by patterns of meal feeding and by any effects of dietary components (e.g. dietary fibre) upon gut transit. Human studies suggest that they will also depend on the initial microbiota composition. There is potential to use the model to explore how the presence of particular functional groups (such as lactate-utilizers (Wang et al 2020)) within an individuals microbiota can influence their gut metabolism and response to dietary intervention. This may indeed be one of the most intriguing and fruitful applications of such modelling approaches in the future.

## Materials and methods

### Software

To facilitate continued research and future model development by other researchers we provide all model code on github (https://github.com/HelenKettle/microPopGut). The R package microPopGut is contained in the file microPopGut 1.0.tar.gz. This can be downloaded and installed in R using install.packages(‘microPopGut 1.0.tar.gz’).

### Microbial Model

The microbial functional group model is based on the model described by Kettle et al. (2015) and implemented using the R package microPop (Kettle et al., 2018). The microbial groups include producers of the three major SCFA detected in fecal samples (acetate, butyrate and propionate) together with utilizers of acetate, lactate, succinate, formate and hydrogen (see Table 1 for a summary, or refer to Kettle et al. (2015) for more detail). The model and its equations are described in detail by Kettle et al. (2015) and Kettle et al. (2018) so only a brief overview is given here The microbial groups are defined as data frames within the R package and these are shown in section 3 of the Supp. Info..

The growth substrates available in the large intestine are divided into four categories: protein (P), non starch polysaccharides (NSP), resistant starch (RS) and sugars (and oligosaccharides and sugar alcohols); for simplicity, all carbohydrate units are regarded as being hexoses. NSP comprise major components of dietary fibre including the structural polysaccharides of the plant cell wall (cellulose, xylan, pectin), whereas RS refers to the fraction of dietary starch that resists digestion in the small intestine. We consider 10 major metabolites that arise from substrate fermentation: acetate, propionate, butyrate, lactate, succinate, formate, hydrogen, carbon dioxide, methane and ethanol. Six of these metabolites (acetate, lactate, succinate, formate, hydrogen and carbon dioxide) are also considered as substrates, because they are known to be consumed by some groups (cross-feeding). It is well known that pH affects growth rate therefore each group is assigned a preferred range of pH within which it can reach its maximum growth rate, but outside of which, its growth is reduced or zero. We model the rate of bacterial growth using Monod kinetics and assume that from 1 g of resource, Y g of biomass is produced. We assume that resource that is taken up by microbes, but not used to produce biomass, is converted to metabolites. If not all of the resource is converted to biomass or to the metabolites represented in our model, it is discarded. This applies, for example, to many diverse fermentation products of proteins (e.g. phenols, amines) that are not among the 10 major products covered by the model. Although the model was initially developed to be run with multiple strains within each functional group, in the current work we do not do this due to the high CPU time associated with multiple compartments.

### Inflow to colon

#### Incoming substrates and water

The main sources of nutrient for microbiota in the colon are complex dietary carbohydrates that are not absorbed higher up the digestive tract. We use a default value of 50 g d^*−*1^ of carbohydrate, C, in our model and we vary the proportion of this which is NSP or RS using the RS fraction (i.e. RS/(RS+NSP) where RS+NSP=C). Based on Cremer et al. (2017) and references therein, about 15 g of bio-available NSP and 30-40 g of RS enter the colon per day which gives us an RS fraction of 0.67-0.9 with average value of 0.78 which we use as our default value. According to Yao et al. (2016) less is known regarding dietary proteins, P, that escape digestion to reach the large intestine, although it is estimated that around 6 - 18 g P reaches the large intestine daily, the majority from the diet and a small proportion from endogenous origins. Given this, here we assume that 10 g d^*−*1^ of undigested P reaches the colon from dietary intake along with a small amount from mucin degradation (approx. 1 g d^*−*1^). Phillips and Giller (1973) state that water enters at approximately 1.5 l d^*−*1^ and about 90% of this is absorbed by the colon. Stephen and Cummings (1980) states that normal fecal daily output in Britain is 100-200 g d^*−*1^ of which 25-50 g d^*−*1^ is solid matter and the rest (50-175 g d^*−*1^) is water. Thus if 90% is absorbed then this indicates water inflow in the range 0.5 - 1.75 l d^*−*1^. The midpoint of this range is 110 g d^*−*1^ of water outflow which, if 90% is absorbed, implies that the inflow of water is approximately 1100 g d^*−*1^. This will clearly vary depending on the host’s oral water intake but we use 1100 g d^*−*1^ as our default value. The default inflow values are summarised in Table 3.

**Table 3:** Summary of default values used in the model. Parameter values for the microbial groups are given in the Supp. Info. (section 3)

| Symbol | Description | Default Value |
| --- | --- | --- |
| $T_t$ | transit time through colon | 1.25 d |
| $\dot{P}_{diet}$ | protein inflow rate | 10 g d<sup>-1</sup> |
| $\dot{C}_{diet}$ | carbohydrate inflow rate | 50 g d<sup>-1</sup> |
| $\dot{W}_{diet}$ | water inflow rate | 1100 g d<sup>-1</sup> |
| $\dot{M}$ | mucin inflow rate | 5 g d<sup>-1</sup> |
| $K_M$ | half saturation constant for Mucin breakdown | 0.5 g l<sup>-1</sup> |
| $a_W$ | rate of water absorption by host | 3 d<sup>-1</sup> |
| $a_Z$ | rate of SCFA absorption by host | 9.6 d<sup>-1</sup> |

#### Meals

The normal human diet does not consist of continuous fixed inflow of substrate; for a more realistic substrate inflow to the colon we simulate eating 3 meals a day with randomly varying composition. We then approximate the passage of these meals through the stomach and small intestine to obtain a smoothed time series for substrate entering the colon. Note that we are not simulating all the food ingested by the host (most of which will not reach the colon) but rather simply trying to produce a more realistic time series for the substrates that we know reach the colon.

We specify three meals per day each with a duration of 30 minutes. This time-series is then passed through a one-compartment ordinary differential equation model representing the time spent in the stomach and small intestine (estimated to take 7 hours), i.e.

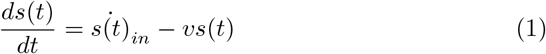

where *v*=3.4 d^*−*1^ (inverse of 7 h transit time in days); 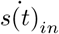 is time series representing 3 meals a day (g d^*−*1^) and *t* is time in days. The inflow to the colon (i.e. the outflow from small instestine) is given by *vs*(*t*). The composition (in terms of P, NSP, RS and water (W)) of these meals varies randomly around the mean of each component (Table 3) for each meal. To generate such random fluctuations we draw samples for each meal from a gamma distribution (since this is always above zero) defined by a scale parameter (*γ*_*s*_) and the daily average inflow of the substrate (g d^*−*1^). We assume the magnitude of the substrate fluctations are proportional to the mean value. Preliminary simulations showed that *γ*_*s*_ equal to half the mean value of each substrate gave a good variation for P, RS and NSP, and for water variation we assumed *γ*_*s*_ was one tenth of the incoming daily flow. The distributions and flow patterns are shown in Fig. 9.

**Figure 9:**
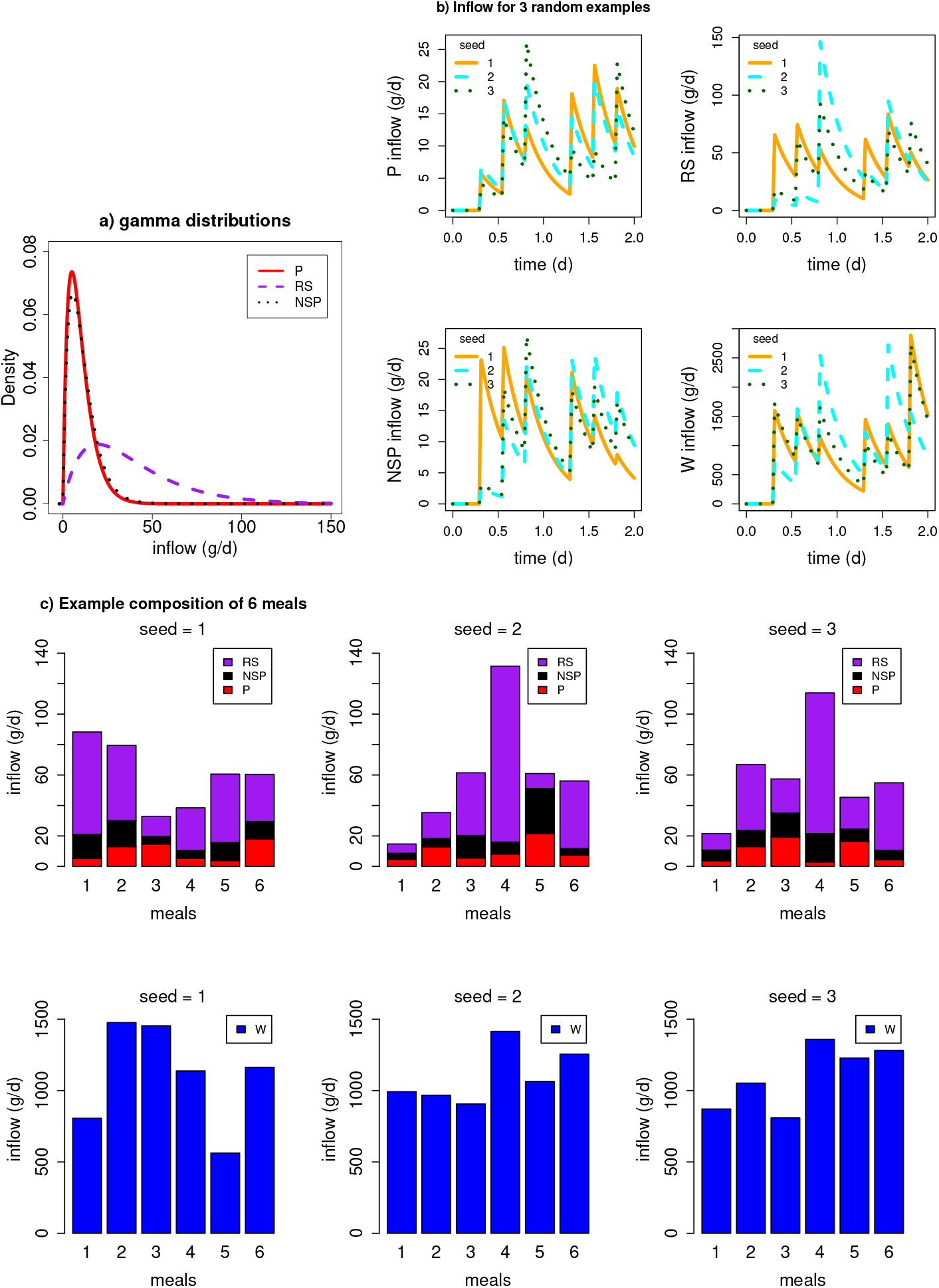
a) Gamma distribution from which random values are drawn to generate the composition of each meal (note water is not shown due to the large difference in magnitude between water and dietary substrates). b) The substrate inflow time series to the proximal colon after passing through the small intestine. Examples shown for 3 stochastic simulations starting with different seeds. c) Barplots showing the composition of 6 meals over 2 days for 3 different stochastic simulations.

#### Mucin

There is a further input of protein and carbohydrate from the host via the breakdown of host-released mucin by many strains in the B group (Ravcheev and Thiele, 2017) and in our NBFD group (Crost et al., 2013). It is estimated that 2.7-7.3 g d^*−*1^ of mucin, denoted 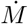, is secreted into the colon (Florin et al., 1991), therefore we take the midpoint value 5 g/d. We assume our mucin degraders break down 1 g of mucin into 0.05 g sulphate, 0.2 g P and 0.75 g C, based on Sung et al. (2017), but consider their yield on mucin to be negligible compared with growth on other substrates. We split C equally between NSP and RS - this arbitrary choice did not affect model results since C from mucin (3.75 g d^*−*1^ maximum) is much less than dietary C (50 g d^*−*1^), but this should be revised if considering very different dietary drivers. Since the compartments of the colon are not equal-sized we assume that the rate of mucin entering the colon is divided through the model compartments proportional to their relative volumes. We assume this enters the colon at a fixed, continuous rate and mucinderived P and C are a function of the mass of mucin degraders, *D*_*M*_ (B and NBFD), such that,

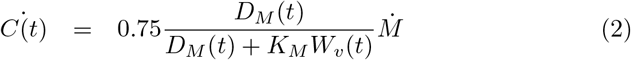

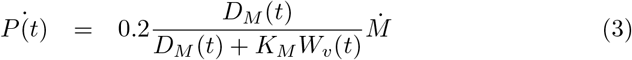

where *C, P, D*_*M*_ are in mass units and the over dot indicates a rate (e.g. g d^*−*1^), *t* is time and *W*_*v*_ is the volume of water in the model compartment. *K*_*M*_ (g l^*−*1^) is chosen such that if *D*_*M*_ << *K*_*M*_ *W*_*v*_ then there is minimal breakdown of mucin and if *D*_*M*_ >> *K*_*M*_ *W*_*v*_ there is maximal breakdown. The smaller the value of *K*_*M*_ the more breakdown there will be at low concentrations of mucin degraders. We set *K*_*M*_ =0.5 g l^*−*1^ based on preliminary model simulations.

### Absorption by host

SCFA and water are both absorbed by the host through the gut wall; over 95% of SCFA (Topping and Clifton, 2001) and approximately 90% of incoming water is absorbed (Phillips and Giller, 1973). Experiments by Ruppin et al. (1980) found that the absorption rates of SCFA to be approximately 0.4 h^*−*1^ (i.e. 9.6 d^*−*1^) with little difference in rates between the different SCFA (Ruppin et al. (1980), Topping and Clifton (2001)).

We can estimate mathematically the specific water absorption rate required to give 90% absorption of inflowing water for a given number of compartments in the colon (*N*) and a given transit time, *T*_*t*_, using

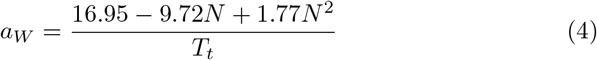

(see Supp. Info. Section 1.3 for the derivation). As a rough estimation, for a 3 compartment model with a transit time 1-1.5 days, gives *a*_*W*_ *≈* 3 d^*−*1^ (Supp. Info. Fig. S1a). Given this will not be significantly affected by the microbial model (microbial uptake/production of water is small) this is a robust estimation.

To estimate the value of the specific absorption rate of SCFA, *a*_*Z*_, we used a simple model (see Supp. Info. sections 1.1 and 1.4). Estimating the value of the specific absorption rate of SCFA based on the values of SCFA given in the verification criteria and given our estimate for *a*_*W*_ we found that it was necessary for the specific absorption rate to change along the colon (see Supp. Info. section 1.4). The best estimates were given by *a*_*Z*_ values of 25.2, 4.2 and 9.2 d^*−*1^ in the proximal, transverse and distal colon respectively. However, in the interests of a robust model (i.e. the fewer parameter values, the better) we made the decision to use one value for *a*_*Z*_. Since the experimental value of 9.6 d^*−*1^ compares well with our estimate in the distal colon we set *a*_*Z*_=9.6 d^*−*1^ throughout. It should be noted though that our model results could potentially be improved by varying *a*_*Z*_ between model compartments.

### pH

Calculating pH in our model is not straightforward due to a lack of necessary state variables as well as pH buffering via secretions from the host. However, observations tell us the pH in the colon goes from 5.7 in the proximal, 6.2 in the transverse and 6.6 in the descending colon and TSCFA in these regions is around 123 mM, 117 mM and 80 mM respectively (Cummings et al., 1987). Therefore an approximate approach is to simply make pH a function of TSCFA.

Fitting a line through the above points gives us the following relationship

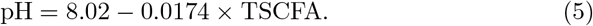

which we further limit by setting the minimum and maximum pH values at 5 and 8 respectively i.e. if the TSCFA values give predicted pH outside of this range (Fig. 10).

**Figure 10:**
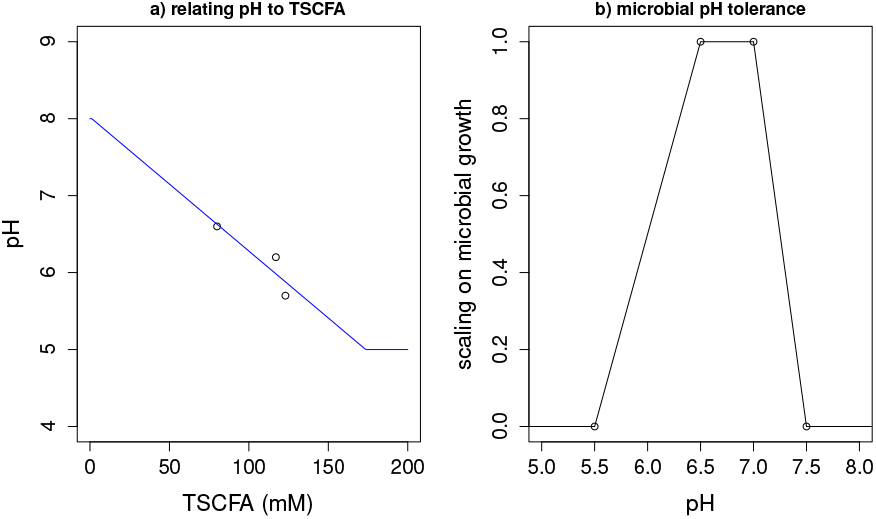
a) Relating pH to TSCFA using Eq. 5 and data from (Cummings et al., 1987). b) Example of microbial tolerance to pH. A pH tolerance function of this form is specified individually for each microbial group in our model.

The impact of pH on microbial growth is modelled via a pH limitation function whereby there is a range over which there is no limit on growth but outside of this range the growth rate decreases linearly to reach zero at the specified outer limits. Thus there are 4 parameters used to describe the pH tolerance – 2 for the inner range where there is no limit on growth and 2 for the outer range outside which there is no growth – an example is shown in Fig. 10. The pH tolerance range for each microbial group is specified under the entry ‘pHcorners’ in the data frame for each group and shown in Supp. Info. section 3.

### Fecal outflow

Fecal outflow (g d^*−*1^) at time, *t*, is given by *m*_*d*_(*t*)*V*_*d*_ where *m*_*d*_(*t*) is the mass in the distal colon (i.e. microbes, unconsumed substrate, microbial metabolites and water) and *V*_*d*_ is the specific wash out rate from the colon (the inverse of the time spent in the distal colon). For continuous outflow (as is used in most gut models) we compute the specific wash out rate from each compartment by assuming the fraction of time spent in compartment, *i*, is proportional to its volume fraction, thus

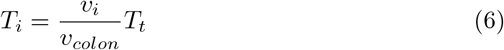

where *v*_*i*_ is the volume of compartment *i* and *v*_*colon*_ is the total volume of the colon. The specific wash out rate is then *V*_*i*_ = 1*/T*_*i*_.

If we introduce bowel movements then, assuming the distal colon is approximately emptied for each bowel movement, the total transit time is given by

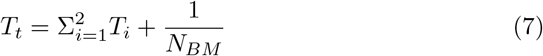

where *N*_*BM*_ is the number of bowel movements per day. For example, using volume measurements (Table 1B) and assuming a total transit time of 1 day would mean about 45% of the transit time is spent in the proximal and transverse colon and about 55% of the day spent in the distal, which would be similar to 2 bowel movements per day. In model experiments where we vary the number of bowel movements per day we also change the time spent in the rest of the colon since we assume increased bowel movements are indicative of a general increase in passage rate. We estimate the wash out rate from the colon during a bowel movement, *V*_*BM*_, by

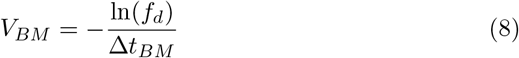

where *f*_*d*_ is the fraction of mass left in the distal colon after the bowel movement and Δ*t*_*BM*_ is the time taken for the bowel movement (d). For example, if a bowel movement takes 10 minutes to remove 90% of the contents of the distal colon then *V*_*BM*_ is 332 d^*−*1^. This is not affected by the number of bowel movements per day.

## Acknowledgments

We thank the Scottish Goverment’s Rural and Environment Science and Analytical Services Division (RESAS) for funding this research.

